# Adaptive landscape flattening allows the design of both enzyme:substrate binding and catalytic power

**DOI:** 10.1101/771824

**Authors:** Vaitea Opuu, Giuliano Nigro, Emmanuelle Schmitt, Yves Mechulam, Thomas Simonson

**Affiliations:** Laboratoire de Biochimie (CNRS UMR7654), Ecole Polytechnique, Palaiseau, France

**Keywords:** computational protein design, Proteus software, aminoacyl-tRNA synthetase, molecular mechanics

## Abstract

Designed enzymes are of fundamental and technological interest. Experimental directed evolution still has significant limitations, and computational approaches are complementary. A designed enzyme should satisfy multiple criteria: stability, substrate binding, transition state binding. Such multi-objective design is computationally challenging. Two recent studies used adaptive importance sampling Monte Carlo to redesign proteins for ligand binding. By first flattening the energy landscape of the apo protein, they obtained positive design for the bound state and negative design for the unbound. We extend the method to the design of an enzyme for specific transition state binding, *i.e.*, for catalytic power. We consider methionyl-tRNA synthetase (MetRS), which attaches methionine (Met) to its cognate tRNA, establishing codon identity. MetRS and other synthetases have been extensively redesigned by experimental directed evolution to accept noncanonical amino acids as substrates, leading to genetic code expansion. We redesigned MetRS computationally to bind several ligands: the Met analog azidonorleucine, methionyl-adenylate (MetAMP), and the activated ligands that form the transition state for MetAMP production. Enzyme mutants known to have azidonorleucine activity were recovered, and mutants predicted to bind MetAMP were characterized experimentally and found to be active. Mutants predicted to have low activation free energies for MetAMP production were found to be active and the predicted reaction rates agreed well with the experimental values. We expect the present method will become the paradigm for computational enzyme design.

## 1 Introduction

One of the most important challenges in computational protein design (CPD) is to modify a protein so that it will bind a given ligand [1–4]. This is essential for problems like enzyme design, biosensor design, and constructing tailored protein assemblies. To design ligand binding means optimizing a free energy difference between bound and unbound states. This two-state optimization is not directly tractable by the most common CPD methods, such as simulated annealing, plain Monte Carlo (MC), or simple branch-and-bound and dead end elimination methods [4, 5]. Rather, most studies have used either heuristic methods that optimize the *bound state* energy [1–4, 6], or enumeration methods that are rigorous but expensive and explore a limited free energy range [7–10].

Recently, a new approach was proposed, using Monte Carlo simulation and importance sampling. The energy landscape in sequence space is flattened adaptively over the course of a simulation, thanks to a bias potential [11]. Flattening can be done for the bound state, the unbound state, or both [12]. Remarkably, this leads to a situation where sequence variants are sampled according to a Boltzmann distribution controlled by the *binding free energy*, exactly the quantity we want to select for. Several variations have been employed, including one that used molecular dynamics instead of MC [13]. The method allows sequences to be designed for binding affinity, but also binding *specificity*. This is especially important for enzyme design, since catalytic power is directly related to the enzyme’s ability to preferentially stabilize the transition state [14]. We apply the method here to an enzyme of biological and technological importance, methionyl-tRNA synthetase. We demonstrate that the method can be used to design an enzyme for its catalytic power.

Each aminoacyl-tRNA synthetase (aaRS) attaches a specific amino acid to a tRNA that carries the corresponding anticodon, establishing the genetic code [15]. Several aaRSs have been engineered experimentally to bind noncanonical amino acids (ncAAs) [16–20]. Obtaining an aaRS that binds an ncAA and uses it as a substrate is a key step to allow the ncAA to become part of an expanded genetic code [17, 20, 21]. The ncAA can then be genetically encoded and incorporated into proteins by the cellular machinery. Several MetRS variants that accepted the ncAA azidonorleucine (AnL) as a substrate were obtained earlier by experimental directed evolution [22]. The AnL azide group can be used for protein labeling and imaging.

The design procedure has two stages. First, a bias potential is optimized adaptively over the course of a MC simulation of the apo protein. The adaptation method is closely analogous to the Wang-Landau and metadynamics approaches [23, 24]. The bias is chosen so that all the allowed residue types achieve comparable probabilities at all mutating positions. This implies that the free energy landscape in sequence space has been flattened, and the bias of each sequence is approximately the opposite of its apo free energy. In the second stage, the holo state is simulated. The bias is included in the energy function, “subtracting out” the apo free energy. Thus, the method achieves positive design for the bound state and negative design for the unbound. The sequences sampled in the second stage are distributed according to their binding free energies, with tight binders exponentially enriched.

In an analogous procedure, a bias potential can be optimized for the protein bound to one ligand, say *L*. Then a complex with another ligand is simulated, say, *L*′, including the bias. The sequences sampled preferentially in the second simulation are those with a strong binding free energy difference between the two ligands, *i.e.*, the most *specific* binders. Importantly, *L*′ can be an activated, transition state ligand, while *L* is the non-activated substrate. In this case, the first simulation flattens the ground state landscape, while the second preferentially samples sequences that stabilize the transition state, relative to the ground state. Thus, the method can be used to select directly for low activation free energies. It is then straightforward to rank the sampled sequences based on their catalytic efficiency, the ratio between the rate constant for the catalytic step, *k*_cat_ and the Michaelis constant *K*_*M*_.

Here, we report CPD calculations that aim to increase the binding of several ligands by MetRS. We first considered AnL. Three residues in the active site were allowed to mutate. The CPD method was tested for its ability to predict the known experimental variants [22]. The top seven experimental variants were visited by the MC simulations, and six were highly ranked among the predicted sequences. If we ranked the CPD sequences according to binding specificity (AnL vs. Met), several experimental variants also had high ranks. We next considered the natural ligand methionyl adenylate (MetAMP). Another set of three residues near the ligand side chain were allowed to mutate. The wildtype sequence was highly ranked by the computational design. 13 other sequences among the top 40 were tested experimentally and all found to be active. The computed binding free energy differences between variants were mostly in good agreement with the experimental values, obtained from kinetic measurements of the enzyme reaction. Next, we predicted MetRS variants that were specifically designed to bind the transition state for the enzymatic reaction Met+ATP → MetAMP+PP_*i*_. The wildtype enzyme was highly ranked among 5832 possible variants, and for 20 variants that were characterized experimentally, the transition state binding free energies from the simulations were in good agreement with the values deduced from the experimental reaction rates. These calculations represent the first time an enzyme is specifically designed to optimize its transition state binding free energy relative to ground state binding, *i.e.*, its catalytic power. We expect the method will become the paradigm for computational design of enzymes.

## 2 Computational methods

### 2.1 Designing for ligand binding

#### Stage 1: adaptive apo simulation

We consider a polypeptide, with or without a bound ligand. Below, we will use a fixed backbone geometry, but the method is valid with a flexible backbone. Side chains can explore a few discrete conformations, or rotamers, and a few selected positions are allowed to mutate. In a first stage, we perform a MC exploration of the protein with no ligand, using the usual Metropolis-Hastings scheme [25–27]. We gradually increment a bias potential until all the side chain types at the mutating positions have roughly equal populations, thus flattening the free energy landscape. We number the mutating positions arbitrarily 1, …, *p*. The bias *E*^*B*^ at time *t* has the form:

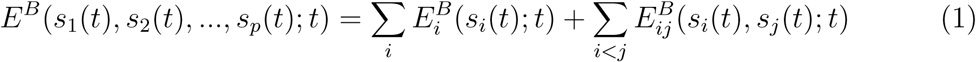

Here, *s*_*i*_(*t*) represents the side chain type at position *i*. The first sum is over single amino acid positions; the second is over pairs. The individual terms are updated at regular intervals of length *T*. At each update, whichever sequence variant (*s*_1_(*t*), *s*_2_(*t*), …, *s*_*p*_(*t*)) is populated is penalized by adding an increment 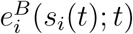 or 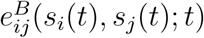 to each corresponding term in the bias. The increments have the form:

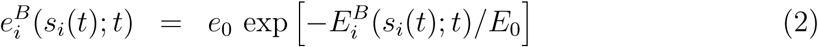

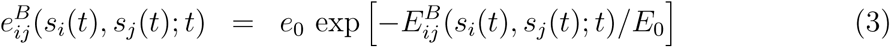

where *e*_0_ and *E*_0_ are constant energies. Thus, the increments decrease exponentially as the bias increases. This scheme is adapted from well-tempered metadynamics [24, 28, 29]. The individual bias terms depend on the system history, and can be written:

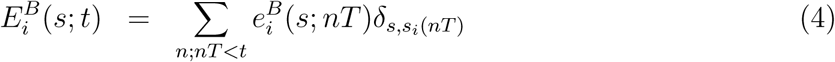

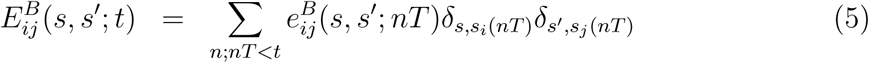

where *δ*_*a,b*_ is the Kronecker delta. Over time, the bias for the most probable states grows until it pushes the system into other regions of sequence space. Two-position biases were implemented in the Proteus software [30, 31] during this work.

#### Stage 2: biased holo simulation

In the second stage, the protein:ligand complex is simulated using the bias potential from stage 1. The sampled population of a sequence *S* is normalized to give a probability, denoted 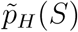, where the subscript means “holo” and the tilde indicates that the bias is present. The apo state probability 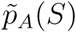 was obtained in stage 1. Both probabilities can be converted into free energies 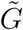:

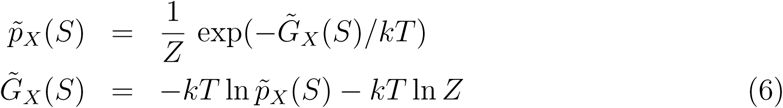

where *X* = *A* or *H* and *Z* is a normalization factor that depends on *X* but not *S*. We also have a relation between the free energies with and without the bias:

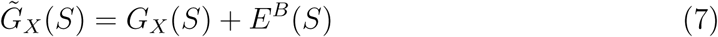

whose (straightforward) derivation is given in Supplementary Material. Note that if the apo state flattening were ideal, 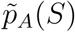 would be a constant, so that (from Eqs. 6, 7) *E*^*B*^(*S*) = −*G*_*A*_(*S*), up to a constant. Thus, the ideal bias is the opposite of the apo free energy.

The binding free energy relative to a reference sequence *R* can be deduced from the populations. We have:

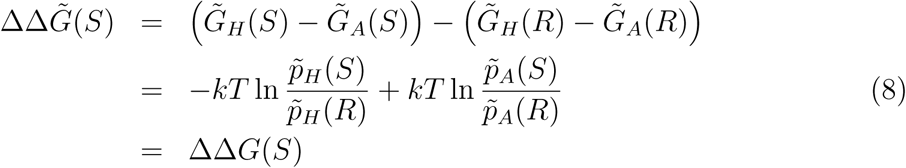

Since the bias is the same in the bound and unbound states, it cancels out from 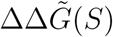, which is equal to the relative binding free energy *in the absence of bias*, ΔΔ*G*(*S*). Although the bias does not appear explicitly in Eq. (8), it is essential for accurate sampling.

In the holo state, the probability of a sequence *S* (with bias) is:

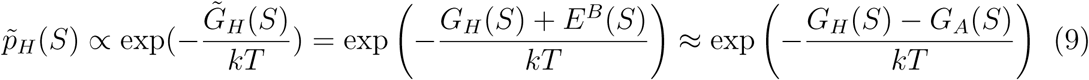

Thus, holo sampling follows a Boltzmann distribution governed by *G*_*H*_(*s*) + *E*^*B*^(*S*), which is approximately the binding free energy *G*_*H*_(*S*) − *G*_*A*_(*S*). This is exactly the quantity we want to design for. If the apo state is well-flattened, the biased holo simulation will be exponentially enriched in tight binders.

### 2.2 Energy function and matrix

The energy was computed using either an MMGBLK or an MMGBSA function (“molecular mechanics + Generalized Born + Lazaridis-Karplus” or “Surface Area”):

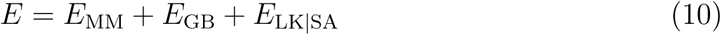

The MM term used the Amber ff99SB force field [30, 32]. The SA term was described earlier [33–35]. The LK term and its parameterization were described earlier [35]. The GB term corresponds to a variant very similar to the one used in Amber, detailed in previous articles [33, 36, 37]. To make the calculation efficient, we compared two strategies. The first used a Native Environment Approximation (NEA), where the GB solvation radii for a given side chain were computed with the rest of the system in its native conformation [36, 38]. The second used a “Fluctuating Dielectric Boundary” (FDB) method, where the GB interaction between two residues *I, J* was expressed as a polynomial function of their solvation radii [39]. These were kept up to date over the course of the MC simulation, so the GB interaction could be deduced with little additional calculation [37, 39]. The solvent dielectric constant was 80; the protein one was 4.0 with the GBSA variants and 6/8 with GBLK [35]. Each solvent model is referred to by its GB variant and nonpolar term; for example, the FDBLK model combines FDB with LK.

To allow very fast MC simulations, we precomputed an energy matrix for each system [34, 40]. For each pair of residues *I, J* and all their allowed types and rotamers, we performed a short energy minimization (15 conjugate gradient steps) [30]. The backbone was fixed (in its crystal geometry) and the energy only included interactions between the two side chains and with the backbone. At the end of the minimization, we computed the interaction energy between the two side chains. Side chain–backbone interaction energies were computed similarly (and formed the matrix diagonal) [30].

### 2.3 Structural models

#### MetRS:AnL and MetRS:MetAMP complexes

For MetRS:AnL, we started from the crystal structure of a complex between a triple mutant of *E. coli* MetRS and AnL (PDB code 3H9B) [41]. The protein mutations were L13S, Y260L, H301L. We refer to this mutant as SLL. The protein backbone and side chains more than 20 Å from the ligand were held fixed. The other side chains were allowed to explore rotamers, taken from the Tuffery library, augmented to allow multiple orientations for certain hydrogen atoms [42, 43]. Side chains 13 and 301 were allowed to mutate into the following 14 types: ACDEGHIKLMQSTV; position 260 was allowed to mutate into the same types, except that Tyr replaced Asp. Thus, there were 14^3^ = 2744 possible sequences in all. Histidine protonation states at non-mutating positions were assigned by visual inspection of the 3D structure. System preparation was done using the protX module of the Proteus design software [31].

For MetRS:MetAMP, we started from a crystal complex (PDB code 1PG0) between *E. coli* MetRS and a methionyl adenylate (MetAMP) analogue [44]. The protein backbone and side chains more than 20 Å from the ligand were held fixed. The other side chains were allowed to explore rotamers [42, 43]. Side chains 13, 256 and 297 were allowed to mutate into all types except Gly or Pro, for a total of 5832 possible sequences in all. Histidine protonation states at non-mutating positions were assigned by visual inspection of the 3D structure.

#### Unfolded state

The unfolded state energy was estimated with a tri-peptide model [45]. For each mutating position, side chain type, and rotamer, we computed the interaction between the side chain and the tri-peptide it forms with the two adjacent backbone and C_*β*_ groups. Then, for each allowed type, we computed the energy of the best rotamer and averaged over mutating positions. The mean energy for each type was taken to be its contribution to the unfolded state energy. The contributions of the mutating positions were summed to give the total unfolded energy.

### 2.4 Ligand force field and rotamers

#### Force field

For the AnL azido group, we used atomic charges and van der Waals parameters obtained earlier for azidophenylalanine [46]. Parameters for the implicit solvent energy terms were assigned by analogy to existing groups. For methionyl adenylate (MetAMP), we mostly used existing Met and AMP parameters. For atoms close to the Met:AMP junction, we used atomic charges computed earlier for ThrAMP (G. Monard, personal communication) from *ab initio* quantum chemistry, in a manner consistent with the rest of the Amber force field [32]. Van der Waals parameters for atoms near the junction were assigned the same types as in Met or AMP. Parameters for bond lengths, angles and dihedrals involving junction atoms were taken from the experimental geometry of MetAMP. Stiffness parameters were assigned by analogy to existing parameters. The complete set of parameters for AnL and MetAMP is in Supplementary Material.

#### Rotamers

AnL was positioned in the protein complex so that its backbone had the position occupied in the MetRS:AnL crystal structure [41]. The ligand’s side chain was allowed to explore rotamers. These were defined by the usual side chain rotamers of Met [42, 43]. We started by positioning Met in the pocket by superimposing it on AnL in the mutant MetRS:AnL crystal complex (PDB code 3H9B). We then positioned the 17 Met side chain rotamers from the Tuffery library. We extracted the AnL side chain from the experimental complex and superimposed it on each of the 17 Met rotamers, producing 17 AnL conformers. Finally, for each one, we performed a short energy minimization with the AnL backbone held fixed. The 17 minimized conformers defined the AnL rotamers. Notice that with this procedure, the azido group always had the same orientation relative to the aliphatic part of the AnL side chain. The AnL rotamers are shown in Supplementary Material. For MetAMP, we allowed the Met rotamers from the Tuffery library, with the rest of the ligand held fixed. The *ϕ* and *ψ* dihedral angles around the MetAMP C_*α*_ were not allowed to rotate and the whole AMP moiety stayed fixed.

### 2.5 Modeling the MetRS transition state complex

A model for the ground state ligands Met + ATP was first obtained, starting from the crystal complex between MetAMP and PP*_i_* (PDB code 3KFL). The covalent structure was reset to that of Met + ATP and the geometry was adjusted by a short energy minimization. The complex included a magnesium ion. Next, a model for the activated ligand [Met:ATP]^‡^ was obtained, starting from the Met + ATP complex. A covalent bond was introduced between the reacting Met carboxylate oxygen and the *α* phosphorus atom. The lengths for this bond and the symmetric one on the other side of the phosphorus were set to 2.4 Å, based on an *ab initio* model of the Tyr + ATP reaction (Thomas Gaillard, personal communication). Planarity restraibts were imposed on the phosphorus and the three *α* phosphate oxygens. A short energy minimization was done. This led to the expected *α* phosphate geometry, with three oxygens in plane and two perpendicular (Fig. 1), as expected for in-line attack of the Met carboxylate on the phosphate [44, 47–49]. *Ab initio* atomic charges were then computed for the entire activated ligand in this geometry, from a Merz-Kollman population analysis of the HF/6-31G* wavefunction [32], using Gaussian 9.0. The magnesium ion, which bridges the *α, β* and *γ* phosphates, was included in the calculation. The resulting charges were applied to atoms close to the *α* phosphate group, while other atoms kept their usual Met or ATP charges. Small manual adjustments were made to establish the correct total charge of −4. The final Mg charge was +1.5. Charges are in Supplementary Material.

**Figure 1:**
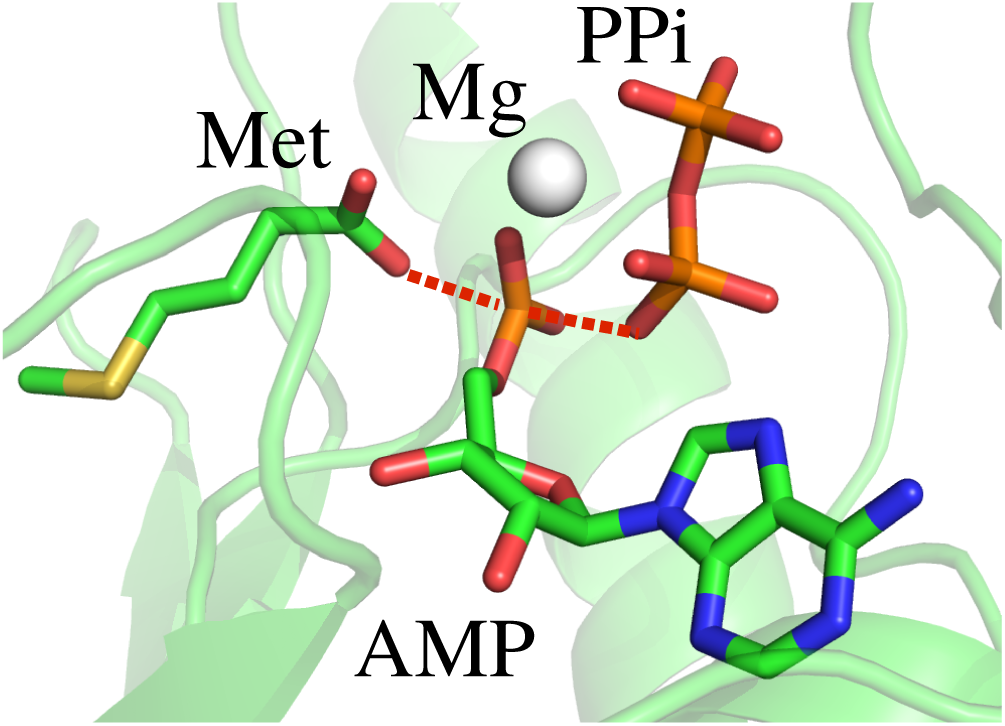
Closeup of the transition state ligands for MetAMP formation.

The geometry of the protein around the ligands was relaxed slightly by performing a short, restrained molecular dynamics simulation, with the ligands held fixed. The entire system was placed in a large box of explicit TIP3P water [50]. Harmonic restraints were applied to nonhydrogen atoms, with force constants that decreased gradually from 5 to 0.5 kcal/mol/Å^2^ over 575 ps of dynamics, performed with the NAMD program [51]. The final protein geometry was used for the design calculations.

### 2.6 Monte Carlo simulations

To optimize the bias potential, we performed MC simulations of the apo state with bias updates every *T* = 1000 steps, with *e*_0_ = 0.2 kcal/mol and *E*_0_ = 50 kcal/mol [12]. During the first 10^8^ MC steps, we optimized a bias potential including only single-position terms. There were *p* = 3 mutating positions, which all contributed to the bias. In the second stage, we ran MC or (in one case: MetAMP complex with the FDBSA solvent model) Replica Exchange MC (REMC) simulations of 5.10^8^ MC steps [27], using 8 replicas with thermal energies (kcal/mol) of 0.17, 0.26, 0.39, 0.59, 0.88, 1.33, 2.0 and 3.0. Temperature swaps were attempted every 500 steps. All the replicas experienced the same bias potential. Both stages used 1- and 2-position moves.

### 2.7 Experimental mutagenesis and kinetic assays

#### Purification of wildtype and mutant MetRS

Throughout this study, we used a His-tagged M547 monomeric version of *E. coli* MetRS, fully active, both in vitro and in vivo [41]. The gene encoding M547 MetRS from pBSM547+ [52, 53] was subcloned into pET15blpa [54] to overproduce the His-tagged enzyme in *E. coli* (Nigro et al., submitted). Site-directed mutations were generated using the QuickChange method [55], and the whole mutated genes verified by DNA sequencing. The enzyme and its variants were produced in BLR(DE3) *E. coli* cells. Transformed cells were grown overnight at 37°C in 0.25 L of TBAI autoinducible medium containing 50 *µ*g/ml ampicillin. They were harvested by centrifugation and resuspended in 20 ml of buffer A (10 mM Hepes-HCl pH 7.0, 3 mM 2-mercaptoethanol, 500 mM NaCl). They were disrupted by sonication (5 min, 0°C), and debris was removed by centrifugation (15,300 G, 15 min). The supernatant was applied on a Talon affinity column (10 ml; Clontech) equilibrated in buffer A. The column was washed with buffer A plus 10 mM imidazole and eluted with 125 mM imidazole in buffer A. Fractions containing tagged MetRS were pooled and diluted ten-fold in 10 mM Hepes-HCl pH 7.0, 10 mM 2-mercaptoethanol (buffer B). These solutions were applied on an ion exchange column Q Hiload (16 mL, GE-Healthcare), equilibrated in buffer B containing 50 mM NaCl. The column was washed with buffer B and eluted with a linear gradient from 5 to 500 mM NaCl in buffer B (2 ml/min, 10 mM/min). Fractions containing tagged MetRS were pooled, dialyzed against a 10 mM Hepes-HCl buffer (pH 7.0) containing 55% glycerol, and stored at −20°C. The homogeneity of the purified MetRS was estimated by SDS-PAGE to be higher than 95%.

#### Measurement of ATP-PPi exchange activity

Prior to activity measurements, MetRS was diluted in a 20 mM Tris-HCl buffer (pH 7.6) containing 0.2 mg/ml bovine serum albumin (Aldrich) if the concentration after dilution was less than 1 *µ*M. Initial rates of ATP-PPi exchange activity were measured at 25°C as described [56]. In brief, the 100 *µ*l reaction mixture contained Tris-HCl (20 mM, pH 7.6), MgCl2 (7 mM), ATP (2 mM), [^32^P]PPi (1800-3700 Bq, 2 mM) and various concentrations (0-16 mM) of the Met amino acid. The exchange reaction was started by adding catalytic amounts of MetRS (20 *µ*l). After quenching the reaction, ^32^P-labeled ATP was adsorbed on charcoal, filtered, and measured by scintillation counting. kcat and K_*M*_ values were derived from iterative nonlinear fits of the theoretical equation to the experimental values using either MC-fit [57] or Origin (Origin Lab).

## 3 Results

### 3.1 Designing MetRS to bind azidonorleucine

As a first test, we searched for MetRS variants with strong azidonorleucine (AnL) binding. Positions 13, 260 and 301 were allowed to mutate, for comparison to the earlier experimental data [22]. 14 types were allowed at each position (see Methods), for a total of 2744 possible sequences. We compared three variants of the solvent model, which gave similar results. The first stage was to optimize a bias potential that flattened the free energy landscape in sequence space for apo MetRS. We used a bias potential including single-position terms only. After the adaptation period, we ran a further simulation of 10^8^ MC steps to determine the biased populations. With the FDBLK solvent model, 2135 sequences were visited at least 1000 times, thanks to the adaptive bias. The second stage was to simulate the MetRS:AnL complex in the presence of the bias. 920 sequences were visited at least 1000 times; all but one were visited 1000 times in both stages. For these, we used the sampled populations to deduce the AnL binding free energy (Eq. 8), relative to the X-ray sequence. The overall computation time for system setup, energy matrix precalculation and both MC stages was about one day (per solvent model). Sequences sampled with and without the bias and ligand are shown in Fig. 2 as logos.

**Figure 2:**
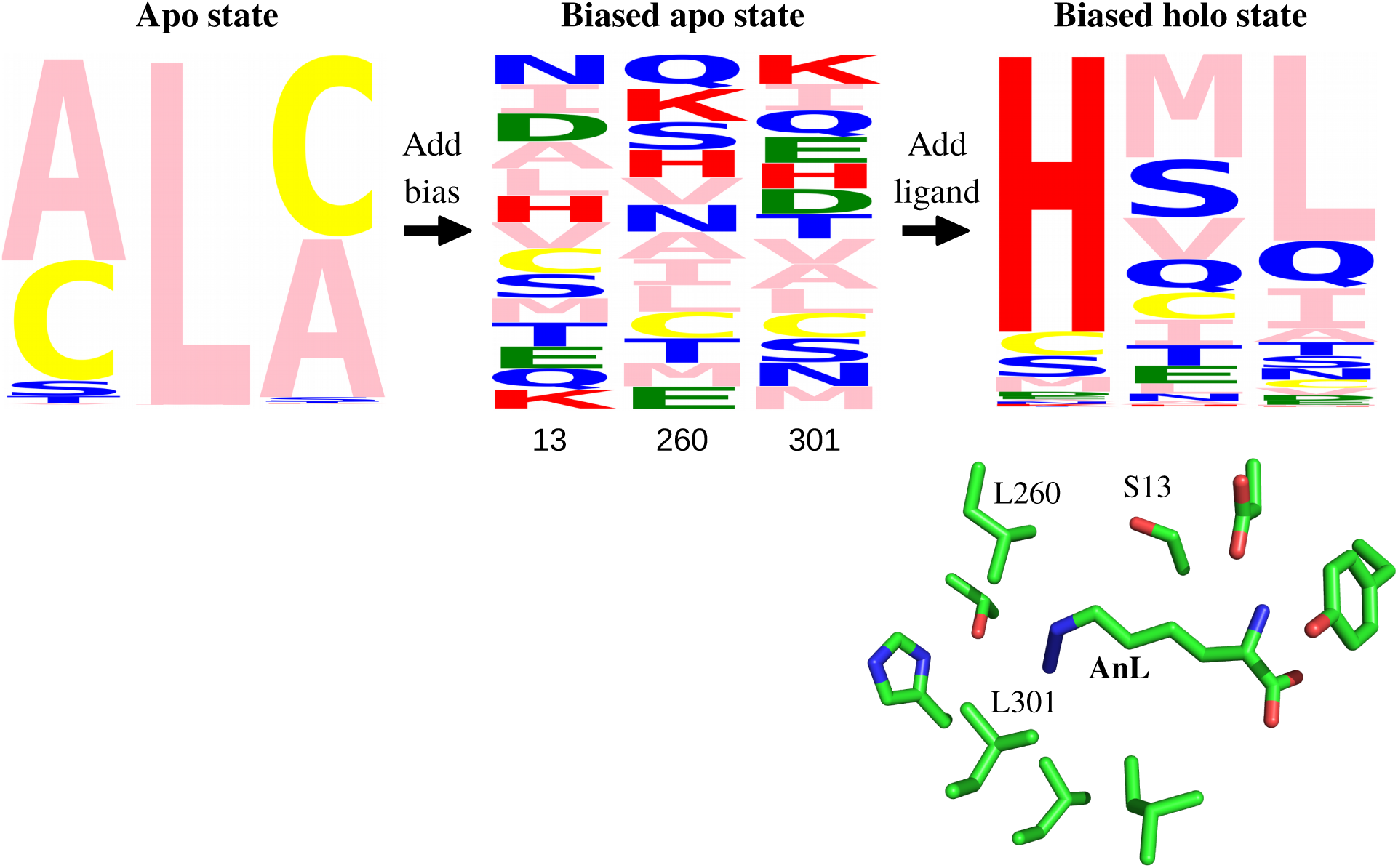
MetRS sequences sampled without and with the AnL ligand (FDBLK solvent model). The three mutating positions, 13, 260, 301 are shown. The logos represent the apo state (left), the biased apo state (middle), and the biased holo state (right). The height of each letter measures the frequency of its type. The 3D view below is a closeup of azidonorleucine (AnL) in the binding pocket, with selected side chains.

Experimental directed evolution had revealed 21 active variants [22], listed in Table 1. Each variant is referred to by the sequence of the three mutating positions; for example, the X-ray variant is SLL. The computational results shown were obtained with the FDBLK variant of the solvent model. The top six experimental sequences were all sampled and had good stabilities and affinities (Table 1). SML was ranked 1st, CVL 7th, SLL (the X-ray variant) 14th and CLL 16th overall. Among the top 20 predictions, there were four variants known to be active. Other predicted variants may also be active, even though they were not revealed by the directed evolution experiments. For the SLL variant, the predicted rotamers for binding site residues were in good agreement with the X-ray structure (Supplementary Material). The FDBSA solvent model gave similar results, while NEASA was slightly poorer (not shown), possibly due to its simpler GB treatment [37].

**Table 1:** MetRS redesigned for AnL binding affinity or specificity

| *a* | *b* | *c* | *d* | *e* | *f* | *g* | *a* | *b* | *c* | *d* | *f* | *e* | *g* |
| --- | --- | --- | --- | --- | --- | --- | --- | --- | --- | --- | --- | --- | --- |
| seq. | pop. | fold | bind | rank | spec. | rank | seq. | pop. | fold | bind | rank | spec. | rank |
| NLL | 62 | 9.1 | 0.9 | 89 | 5.6 | 14 | HTM | 1 | 11.1 | 1.7 | 202 | 8.0 | 118 |
| SLL | 12 | 0.0 | 0.0 | 14 | 0.0 | 1 | LCL | 1 | 20.6 | 4.5 | – | 8.0 | – |
| SML | 4 | 3.9 | -0.6 | 1 | 8.0 | 719 | MAL | 1 | 16.8 | 0.7 | – | 8.0 | – |
| AVL | 3 | 5.9 | 1.2 | 127 | 8.0 | 703 | MSL | 1 | 20.4 | 0.3 | – | 8.0 | – |
| AQL | 2 | 5.4 | 1.6 | 196 | 5.3 | 13 | NVL | 1 | 16.7 | 1.0 | – | 8.0 | – |
| CLL | 2 | -1.2 | 0.0 | 16 | 0.9 | 2 | SCM | 1 | -1.9 | 3.1 | 579 | 8.0 | 669 |
| ACL | 1 | 3.6 | 1.7 | 214 | 8.0 | 667 | SLV | 1 | -2.4 | 1.6 | 196 | 8.0 | 390 |
| CVL | 1 | 6.3 | -0.2 | 7 | 8.0 | 759 | SNL | 1 | 9.7 | 0.3 | 38 | 8.0 | 54 |
| DLI | 1 | 11.8 | 2.0 | 303 | 8.0 | 291 | SSL | 1 | 7.3 | 0.3 | 40 | 8.0 | 310 |
| HCL | 1 | 16.1 | -1.4 | – | 8.0 | – | STL | 1 | 8.0 | 0.6 | 64 | 8.0 | 185 |
|  |  |  |  |  |  |  | FVL | 1 |  |  | unsampled |  |  |

<sup>a</sup>Sequence at the designed positions 13, 260, 301, ranked by <sup>b</sup>population among the experimental clones. <sup>c</sup>Folding and <sup>d</sup>binding free energies (kcal/mol) relative to the X-ray sequence SLL. <sup>e</sup>Rank based on affinity or <sup>g</sup>specificity. <sup>f</sup>Specificity, defined by the binding free energy difference between AnL and Met (relative to SLL). Calculations used the FDBLK solvent.

We also searched for MetRS variants that maximized the AnL binding *specificity*, relative to Met. A bias potential was adaptively optimized for the MetRS:Met complex, then used in a simulation of the MetRS:AnL complex. The mutating positions and allowed types were the same as above. Specificity ranks are included in Table 1. Four of the top six experimental variants had high specificity ranks. The top experimental variant NLL was 14th, the next-best experimental variant SLL was 1st, AQL was 13th, and CLL was 16th. Evidently, selecting for specificity can help reveal active variants.

### 3.2 Redesigning MetRS to bind MetAMP

As a second test, we searched for MetRS variants with a high affinity for the natural ligand methionyl adenylate (MetAMP). These should include the wildtype (WT) sequence and close homologs. Three positions close to the Met side chain (Fig. 3), 13, 256, and 297 were allowed to mutate into all types except Gly or Pro, for a total of 5832 possible sequences. The first stage was to optimize a bias potential that flattened the free energy landscape in sequence space for apo MetRS. We performed calculations with both the FDBSA and the FDBLK variants of the solvent model, which gave similar results. We report the FDBSA results, since they were obtained first and were the basis for choosing which sequences to test experimentally. Selected FDBLK results are also reported. With the FDBSA solvent model, using Replica Exchange MC, 4178 variants were visited at least 1000 times.

**Figure 3:**
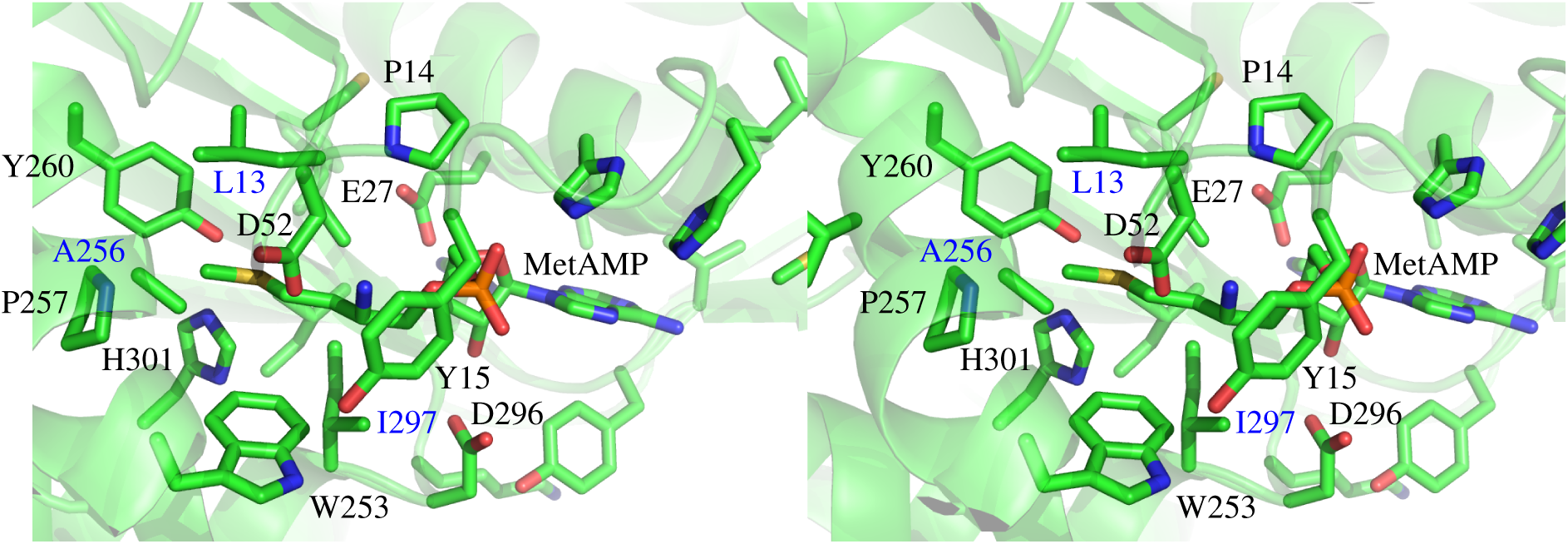
MetRS:MetAMP complex, binding site closeup (stereo). Mutating side chains are 13, 256, 297.

The second stage was to simulate the MetRS:MetAMP complex in the presence of the bias potential. For sequences visited at least 1000 times in both stages (528 sequences), we used the sampled populations to deduce the MetAMP binding free energy (Eq. 8), relative to the wildtype (WT) sequence LAI. The folding energy of each variant was also estimated (see Methods) and sequences less stable than WT by 5 kcal/mol or more were discarded. The top 20 remaining sequences, with the largest binding free energies, are shown in Table 2. The top sequence, CDV, had an Asp at position 256, positioned to form a salt bridge with the MetAMP ammonium group. Its binding free energy, relative to WT, was −1.4 kcal/mol. The next 19 variants had types similar to WT. Their computed binding free energies were close to WT, with relative values between −0.2 and 0.6 kcal/mol. The WT sequence was sixth overall. Among the top 40 variants, 14 mutants were produced experimentally. They were representative of the computational variants, while providing ease of construction (see Methods). CDV was left out, as the A256D mutation, selected for binding, might reduce the catalytic activity. All 14 tested variants had detectable activity, a 100% success rate for the design procedure. One other sequence, SAI, was tested experimentally and found to be active, but did not show up in the MC simulation. Thus the method produced one false negative along with 14 true positives.

**Table 2:**
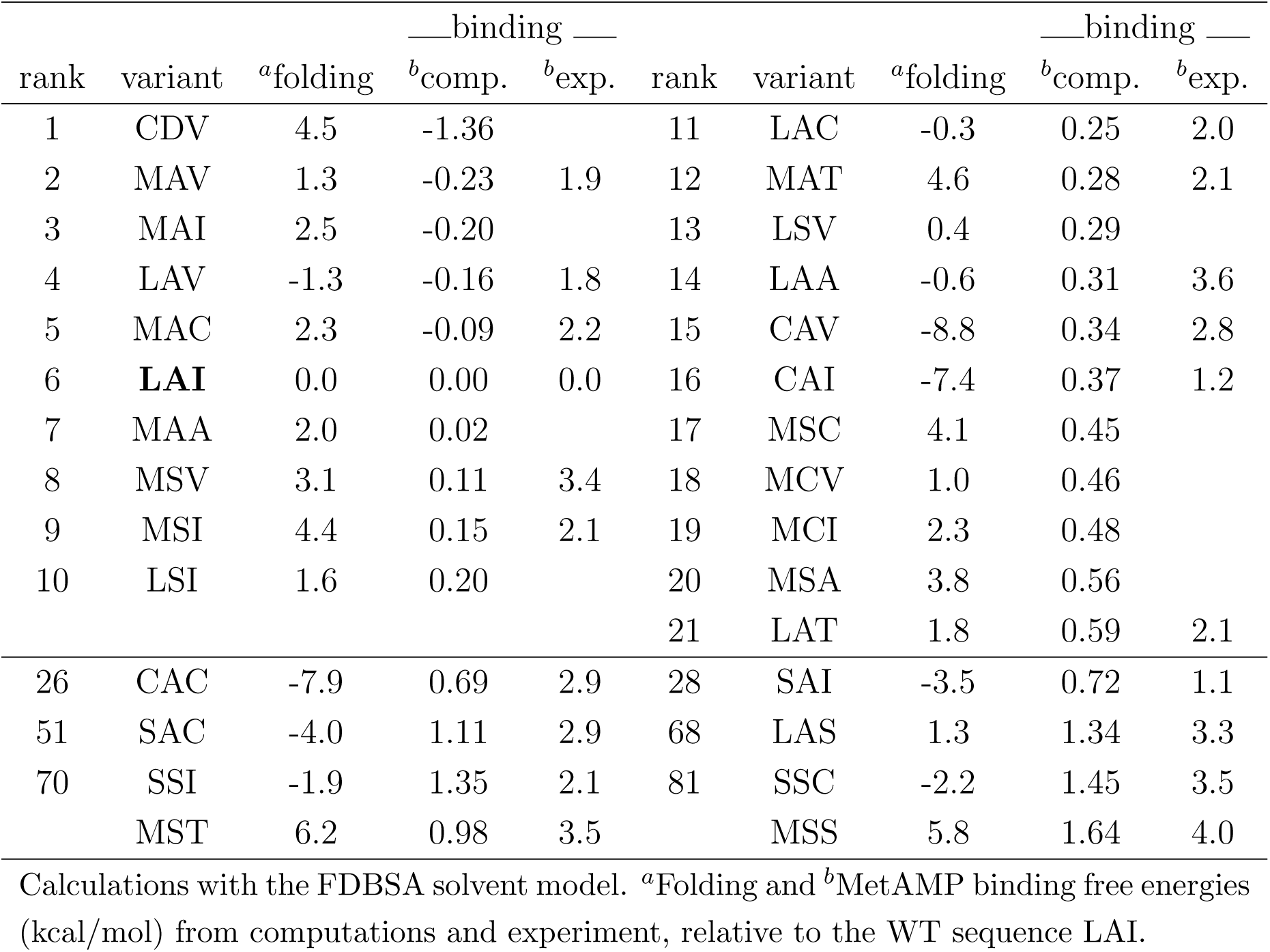
MetRS redesigned for MetAMP binding by mutating positions 13, 256, 297

Going further, we made a quantitative comparison between the computed and experimental binding free energies. The experimental dissociation constants were estimated from the Michaelis constants K_*M*_. In the experimental conditions (excess ATP) and under the usual Michaelis-Menten assumptions [14, 58], *K*_*M*_ represents the dissociation constant for Met binding in the presence of bound ATP. Here, we computed relative binding free energies for binding MetAMP, not Met. Nevertheless, we expect that these MetAMP binding free energy changes can be compared to the experimental Met binding free energy changes; *i.e.*, we make the additional assumption that the relative effects of the mutations will be conserved going from MetAMP to Met+ATP.

Certain mutations at position 297 involved significant changes in the side chain volume, where the largest type, Ile or the smallest type, Ala was introduced or removed. For these, the computed binding free energies departed significantly from the experimental ones. However, if these two types were excluded, there were 27 point mutations between experimental variants, and for these, agreement was very good. The computed binding free energy differences had an rms error of just 0.52 kcal/mol and a mean unsigned error (mue) of 0.42 kcal/mol. The correlation between the experimental and computed sets was 0.59. Fig. 4 shows the binding free energy changes. Note that the good agreement supports the assumption that the experimental *K*_*M*_ values are good proxies for the relative MetAMP binding free energies.

**Figure 4:**
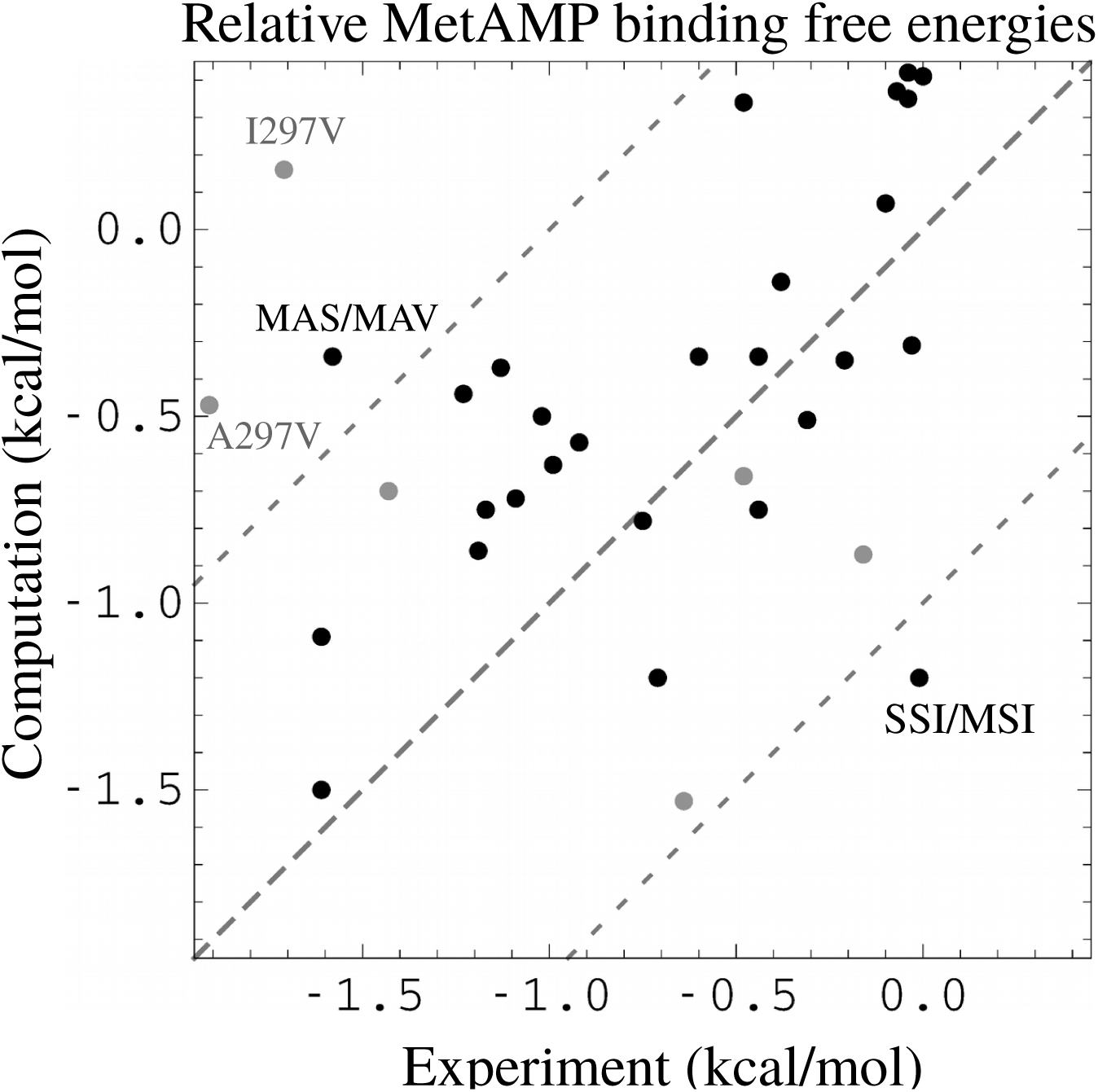
MetRS:MetAMP binding free energies (kcal/mol), relative to the wildtype protein (WT). Shown are data for 27 point mutations. Gray points correspond to two mutations at position 297 that change the side chain volume, and four mutations involving variants that were predicted to be weakly stable (above our 5 kcal/mol threshold, see text) but were produced and measured experimentally nevertheless. Two other mutations with sizable errors are labeled.

With the FDBLK solvent model, results were similar. The WT variant was ranked slightly lower, 20th. The top sequence was SAN, with a binding free energy of −1.3 kcal/mol relative to the WT. 7 of the 14 experimental sequences were ranked among the top 20 predictions. The computed and experimental binding free energy changes associated with point mutations are shown in Supplementary Material. Excluding (as above) mutations involving the types Ile or Ala at position 297, the mue and rms error were 0.76 and 0.98 kcal/mol, respectively, only slightly larger than with FDBSA.

### 3.3 Redesigning MetRS for catalytic power

For enzyme design, it is of great interest to select for a low activation free energy [14]. Therefore, we considered a model of the transition state complex (Fig. 1). The ATP *α* phosphorus was bound to five oxygens: three coplanar and two perpendicular, corresponding to in-line attack of the Met carboxylate. In the first stage, we simulated a competing, ground state complex between MetRS, Met, and ATP. The same three binding pocket residues as above, 13, 256, and 297 were allowed to mutate into all types except Gly, Pro. We optimized a bias potential during the MC simulation, flattening the free energy surface in sequence space. In the second stage, we simulated the transition state complex, with the bias included. All the variants that had been tested experimentally (Table 2) were sampled (WT and 14 variants, plus 5 others that Proteus had predicted to be above our 5 kcal/mol instability threshold). For each one, from the sampled populations, we deduced the free energy difference (Eq. 8) between its ground state and transition state complexes, *i.e.*, its activation free energy. From transition state theory [14], this difference can be identified with the log of the catalytic reaction rate, *k*_cat_. We also computed the Met dissociation free energies for the ground state complexes, which can be identified with the Met Michaelis constants, *K*_*M*_. We first simulated the ground state complex with ATP but no Met, flattening its free energy surface with an adaptive bias. We then simulated the MetRS+Met+ATP complex, including the bias. From the sampled populations, we deduced the Met binding free energy of each variant, relative to WT (Eq. 8). The overall protocol is schematized in Fig. 5.

**Figure 5:**
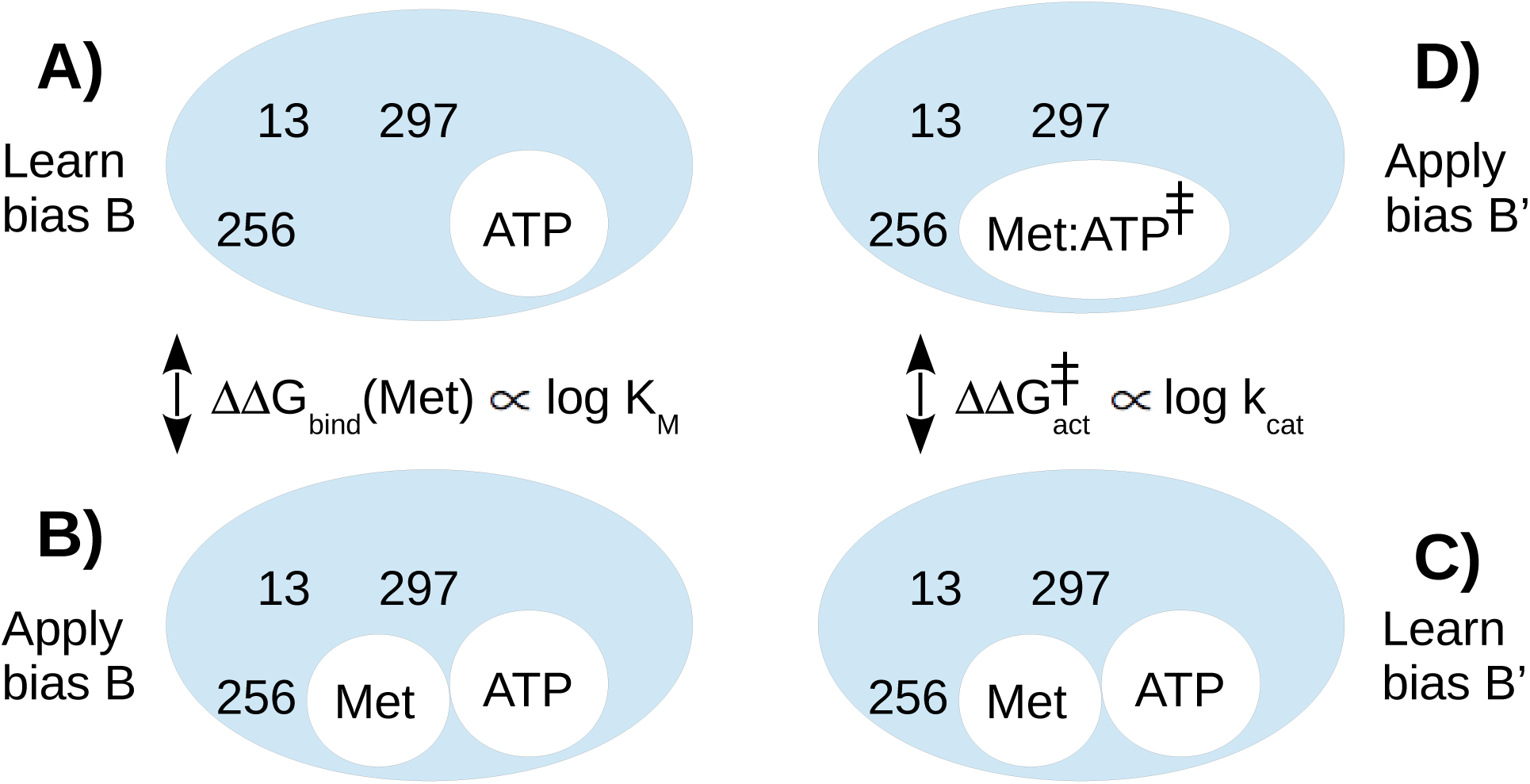
Scheme of the computational method to obtain the catalytic efficiencies *k*_cat_*/K*_*M*_. **A)** A bias *B* is optimized to flatten the sequence landscape of the enzyme without the Met ligand. Mutating positions are 13, 256, 297. **B)** The same bias *B* is used to simulate the complex including Met. Sequences are populated according to their Met binding affinities. **C)** A bias *B*′ is optimized to flatten the sequence landscape of the complex including Met. **D)** *B*′ is used to simulate the transition state complex. Sequences are populated according to their activation free energies. The lefthand simulations yield the predicted *K*_*M*_ values. The righthand simulations yield the predicted *k*_cat_ values.

Fig. 6 compares the *k*_cat_*/K*_*M*_ ratios from experiment and simulations. We refer to them as catalytic efficiencies. We recall that they represent the 2nd order rate constant for the reaction of Met with the MetRS:ATP complex. Fig. 6 shows the quantities *kT* log (*k*_cat_*/K*_*M*_) */* (*k*_cat_*/K*_*M*_)*WT*, which express the catalytic efficiencies on a log scale, in thermal energy units, relative to the WT value. The figure includes WT and 19 other experimental variants. 5 of these had low predicted stabilities and are shown as gray points. WT defines the origin. For the other 14 points, agreement between calculations and experiment is quite good, with a correlation of 0.73 and mean errors of 1.36 kcal/mol (rms) and 1.18 kcal/mol (mue). Experimentally, WT has the largest efficiency. Computationally, two variants are predicted to be slightly better, by 0.9 and 0.2 kcal/mol, respectively, which is less than the mean error. Overall, by designing directly for a low activation free energy, we retrieve all the experimental variants and reproduce the catalytic efficiencies semi-quantitatively.

**Figure 6:**
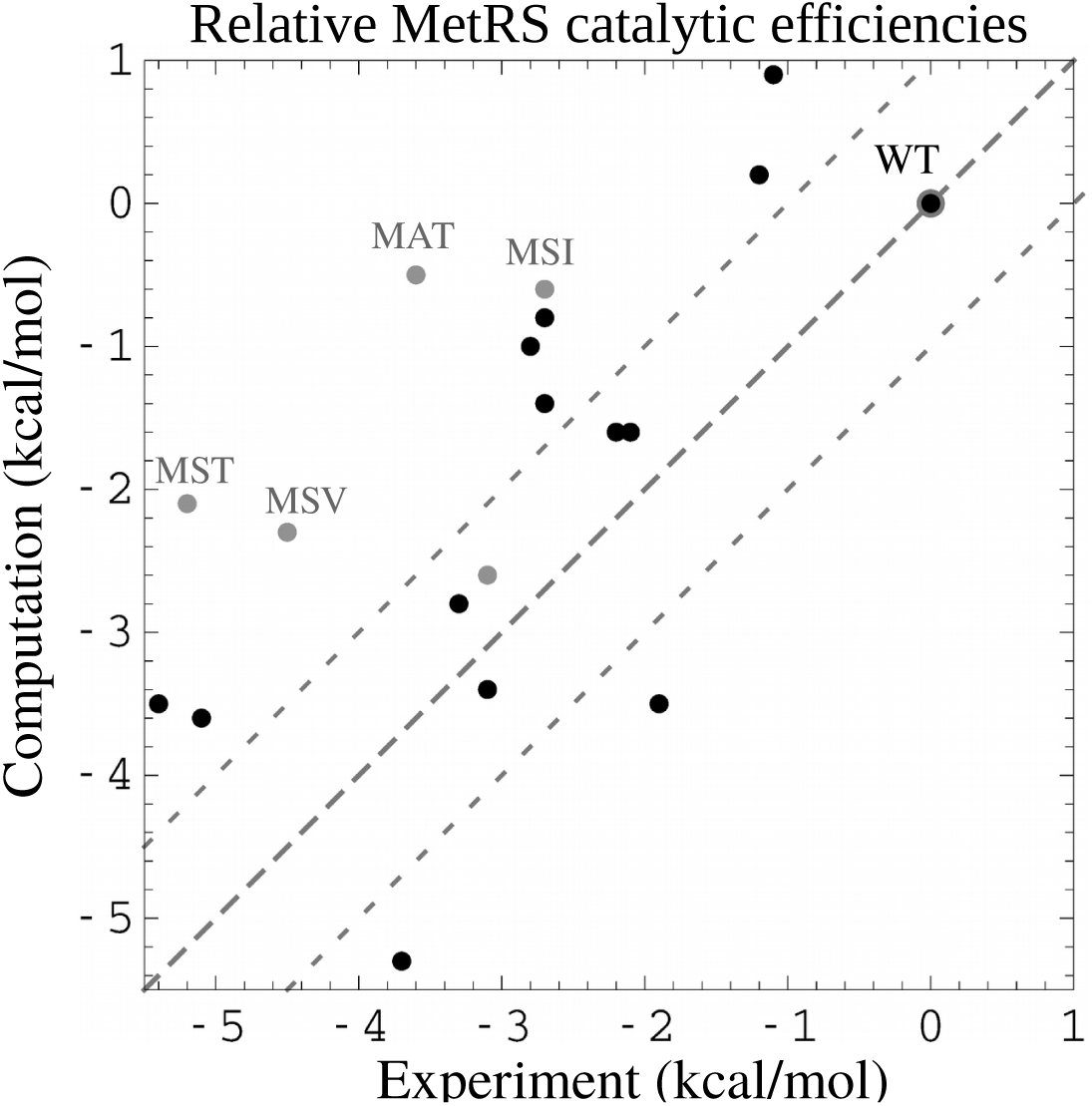
MetRS catalytic efficiencies *kT* log (*k*_cat_*/K*_*M*_) */* (*k*_cat_*/K*_*M*_)*WT* relative to the wildtype (kcal/mol). Four gray points correspond to variants that were predicted to be weakly stable but were produced and measured experimentally nevertheless.

## 4 Discussion

Adaptive importance sampling solves the design problem for ligand binding and specificity. It applies positive design to one state (say, bound) and negative design to the other (unbound). It provides quantitative values for relative binding free energies or activation free energies. Variants sampled for one criterion, such as activation free energy (*k*_cat_), can be reranked *a posteriori* based on another criterion, such as *k*_cat_/*K*_*M*_. *A posteriori* reranking or filtering does not leave out any important solutions; rather, the initial selection brings in too many solutions (*e.g.*, unstable variants), which are then filtered out at very little cost. In the first stage of the procedure, the sampling is very aggressive, if not exhaustive. In the second stage, it does not need to be exhaustive, since the best designs are exponentially enriched, and the unsampled variants are the ones with poor affinities or specificities. If one wants to reveal weak binders or perform reranking on another property, one can also flatten the energy landscape in the second stage. One can also use a more aggressive bias in one or both stages, including two-position biases. Replica Exchange MC can also be used to increase sampling. Using plain MC, one-position biases and no flattening of the holo state, our simulations produced 200 MetRS variants, enriched in tight binders, spanning a 7–8 kcal/mol range of binding free energies.

A difficulty when designing ligand binding is to choose one or more poses for the ligand. Here, we redesigned MetRS in cases where the ligand pose was known from an X-ray structure for one sequence: the SLL sequence in the AnL case and the wildtype sequence in the MetAMP case. For these ligands, we used the experimental ligand pose and protein backbone conformation. Three residues close to the ligand were then allowed to mutate. Not surprisingly, the calculations produced designed sequences that were homologous to the X-ray sequence. The experimental binding free energies in the MetAMP case were well-reproduced (the AnL values are not known). It is likely that other poses exist that would be compatible with other mutations, and would possibly lead to stronger binding. The exploration of such alternate poses was left aside in this work. For the transition state complex, the position of the ligands could also be inferred with some confidence, since the enzyme achieves catalysis with little reorganization or motion of the substrates [15], and X-ray structures were available for the reactant and product complexes. By designing the protein to stabilize this ligand pose, we probably biased the results towards native-like solutions. Here, too, the experimental relative activation free energies were well-reproduced, supporting the structural model.

Overall, agreement with experiment was very good for three MetRS redesign test problems: redesign to bind the AnL ncAA, redesign to bind the natural intermediate MetAMP, and redesign for catalytic power for the reaction that produces MetAMP. Except for the earlier AnL data [22], the experiments were done in this work. Transition state modeling was done simply, by combining two X-ray structures and running a standard quantum chemistry protocol for atomic charges, consistent with the usual Amber force field [32]. Several variants of the implicit solvent model were compared; somewhat better results were obtained using a sophisticated GB approach (FDB variant [37]) that captures the many-body character of polar solvation. All the procedures were carried out with the Proteus software, which is freely available to academics (https://proteus.polytechnique.fr). An entire calculation (setup, matrix calculation, MC simulations, postprocessing) lasted around one day on a 16-core desktop computer. We expect the present adaptive MC method will become the paradigm for computational enzyme design in the future.

## Acknowledgements

We thank Christine Lazennec-Schurdevin for technical assistance, Alexandrine Daniel for performing preliminary MetRS:AnL calculations, Francesco Villa and David Mignon for many helpful discussions, and Thomas Gaillard for discussions on the TyrRS transition state structure.

